# Metabolic intervention by low carbohydrate diet suppresses the onset and progression of neuroendocrine tumors

**DOI:** 10.1101/2022.10.21.507065

**Authors:** Yu Chen, Tatsuki Yamamoto, Yura Takahashi, Tomoka Moro, Tomoko Tajima, Yukiko Sakaguchi, Naoaki Sakata, Akihiko Yokoyama, Susumu Hijioka, Akane Sada, Yuko Tabata, Rieko Ohki

**Affiliations:** Laboratory of Fundamental Oncology, National Cancer Center Research Institute, Tsukiji 5-1-1, Chuo-ku, Tokyo 104-0045, Japan; Department of Regenerative Medicine and Transplantation, Faculty of Medicine, Fukuoka University, Nanakuma 7-45-1, Jonan-ku, Fukuoka, 814-0180, Japan; Tsuruoka Metabolomics Laboratory, National Cancer Center, Yamagata, 997-0052, Japan; Department of Hepatobiliary and Pancreatic Oncology, National Cancer Center Hospital, Tokyo 104-0045, Japan

**Keywords:** neuroendocrine tumor, non-functional pancreatic neuroendocrine tumor, MEN1, ketogenic diet, pituitary neuroendocrine tumor

## Abstract

Insulin signaling often plays a role in the regulation of cancer including tumor initiation, progression, and response to treatment. In addition, the insulin-regulated PI3K-Akt-mTOR pathway plays an important role in the regulation of islet cell proliferation and this pathway is hyperactivated in human non-functional pancreatic neuroendocrine tumors (PanNETs). We therefore investigated the effect of a very low carbohydrate diet (ketogenic diet) on a mouse model that develops non-functional PanNETs to ask how reduced PI3K-Akt-mTOR signaling might affect the development and progression of non-functional PanNET. We found that this dietary intervention resulted in lower PI3K-Akt-mTOR signaling in islet cells and a significant reduction in PanNET formation and progression. We also found that this treatment had a significant effect on the suppression of pituitary NET development. Furthermore, we found that non-functional PanNET patients with lower blood glucose levels tend to have a better prognosis than patients with higher blood glucose levels. This preclinical study shows that a dietary intervention that results in lower serum insulin levels leads to lower insulin signal within the neuroendocrine cells and has a striking suppressive effect on the development and progression of both pancreatic and pituitary NETs.

## Introduction

The pancreas serves two distinctly different functions as an exocrine organ and endocrine organ. As an endocrine organ, pancreatic islet cells release hormones and polypeptides, including insulin and glucagon, that regulate blood sugar levels and multiple other functions. When these pancreatic islet cells become cancerous, they are termed pancreatic neuroendocrine tumors (PanNETs). PanNETs are relatively rare tumors, comprising 1-2% of all pancreatic neoplasms and with an incidence of 0.43 per 100,000; however, this rate has more than doubled in the last 20-30 years ^1^. The majority of PanNETs arise sporadically, but approximately 10% are associated with an underlying genetic syndrome ^2^. PanNETs are divided into functional versus non-functional tumors with about 90% being classified as non-functional. Lesions should be surgically resected when possible, as this is the only potentially curative therapy. Liver metastasis is the most significant prognosis factor in the PanNET progression setting. Most patients, however, that have the metastatic disease are not candidates for resection. The majority of the drugs approved for the management of PanNET show little objective response, providing little progression-free survival or ability to shrink tumors, and thus PanNETs represent a serious unmet medical need in the clinic ^3 4^.

Some PanNETs arise due to a genetic disease called Multiple endocrine neoplasia type 1 (MEN1), an autosomal dominantly inherited tumor syndrome caused by germline mutations of the tumor suppressor gene *MEN1* ^5^. PanNETs are the most prevalent NETs found in MEN1 patients, who will also develop NETs in other organs such as the pituitary; however, the development of a PanNET is a poor prognostic factor. Roughly 50% of patients are diagnosed with PanNETs under age 50, and since most patients have multiple non-functional PanNETs, surgery is not a therapeutic option ^3^. The association between the loss of the *MEN1* gene function and the development of NETs has been confirmed by analyzing *Men1*-deficient mice ^6^. The *Men1*^f/f^-RipCre^+^ mouse in which *Men1* is ablated in pancreatic β-cells and pituitary cells is the most well-characterized MEN1 mouse model ^7, 8^. These mice develop PanNETs and pituitary NETs and their use has contributed to a better understanding of these diseases in vivo. There are several independently derived β cell and pituitary cell-specific *Men1* knock-out mice that are similar but slightly different from one another ^6, 9^. For example, β cell and pituitary cell-specific *Men1*-deficient mice that lack exons 3-8 develop PanNET by 15 months of age, while mice that lack exon 3 develop PanNET by 6 months ^7, 8^.

Insulin signaling plays a role in cancer including tumor initiation, progression and response to treatment ^10^. Studies of some common cancers, such as breast and pancreatic cancers, have shown that hyperinsulinemia enhances tumor development in both humans and mice, while reduced insulin levels reduces tumor development ^11 12 13 14^. Several preclinical studies in mice have shown that dietary interventions leading to low insulin levels in the blood can enhance anticancer therapy and improve the outcomes of several cancers ^15 16^. It is of note that a diet that reduces the serum insulin levels, i.e., a low-carbohydrate ketogenic diet, is safe and has no adverse effects in mice ^17^.

Pancreatic islets are exposed to high levels of insulin, which is released from β cells within the islets^18^. In addition, it has been shown that the insulin-regulated PI3K-Akt-mTOR pathway plays an important role in the regulation of islet cell proliferation and this pathway is hyperactivated in human PanNETs ^19 20 21^. We have previously shown that hyperactivation of the PI3K-Akt-mTOR pathway caused by suppression of the tumor suppressor gene *PHLDA3*, which encodes a repressor of Akt, contributes to the formation of hyperplastic islets in mouse models and to the progression of human PanNETs ^20 21 22 23 24^. These observations led us to investigate the effects of an insulin-lowering diet on PanNET initiation and progression using a newly established non-functional PanNET mouse model, which we fed with a very low carbohydrate diet (ketogenic diet). When fed a normal diet, these mice develop non-functional PanNET by the age of 45 weeks, however when fed a very low carbohydrate diet both initiation and progression of non-functional PanNET were significantly suppressed. We also found that pituitary NET development in these mice is attenuated by a ketogenic diet. We further showed that PanNET patients with higher blood glucose levels tend to have a poorer prognosis than those with lower blood glucose levels. Collectively, these results suggest a novel therapeutic and preventive approach of using a ketogenic diet to treat NET patients, including sporadic and familial non-functional PanNET and pituitary NET patients. This treatment has shown no adverse effects and provides a novel method to fulfill the unmet medical needs of non-functional PanNET and pituitary NET patients.

## Results

### β cell-specific *Men1* knock-out mice that lack the exons 3-6 develop non-functional PanNETs

We made a β cell-specific *Men1* conditional knock-out mouse line by crossing *Men1* exons 3-6 floxed mice with RIP-Cre mice (*Men1*^f/f^-RipCre^+^) ^25^. Similar to previous reports, nearly all of these mice developed PanNET by 10 months of age (Fig. 1A). As shown in Figs. 1B-1D, we found that the body weights, blood glucose levels and serum insulin levels in these mice are similar to those of wild-type mice at the same age. However, insulin expression is reduced in the islets of the *Men1*^f/f^-RipCre^+^ mice at 10 months compared to the wild-type mice at the same age (Fig. 1E). It was previously reported that β cell-specific, *Men1*-deficient mice that lack exons 3-8 or exon 3 develop functional insulinomas with significantly elevated levels of serum insulin levels ^7, 8^. However, we found that the β cell-specific *Men1*-deficient mice we established develop non-functional PanNETs that resemble those found in human MEN1 patients.

**Fig. 1.**
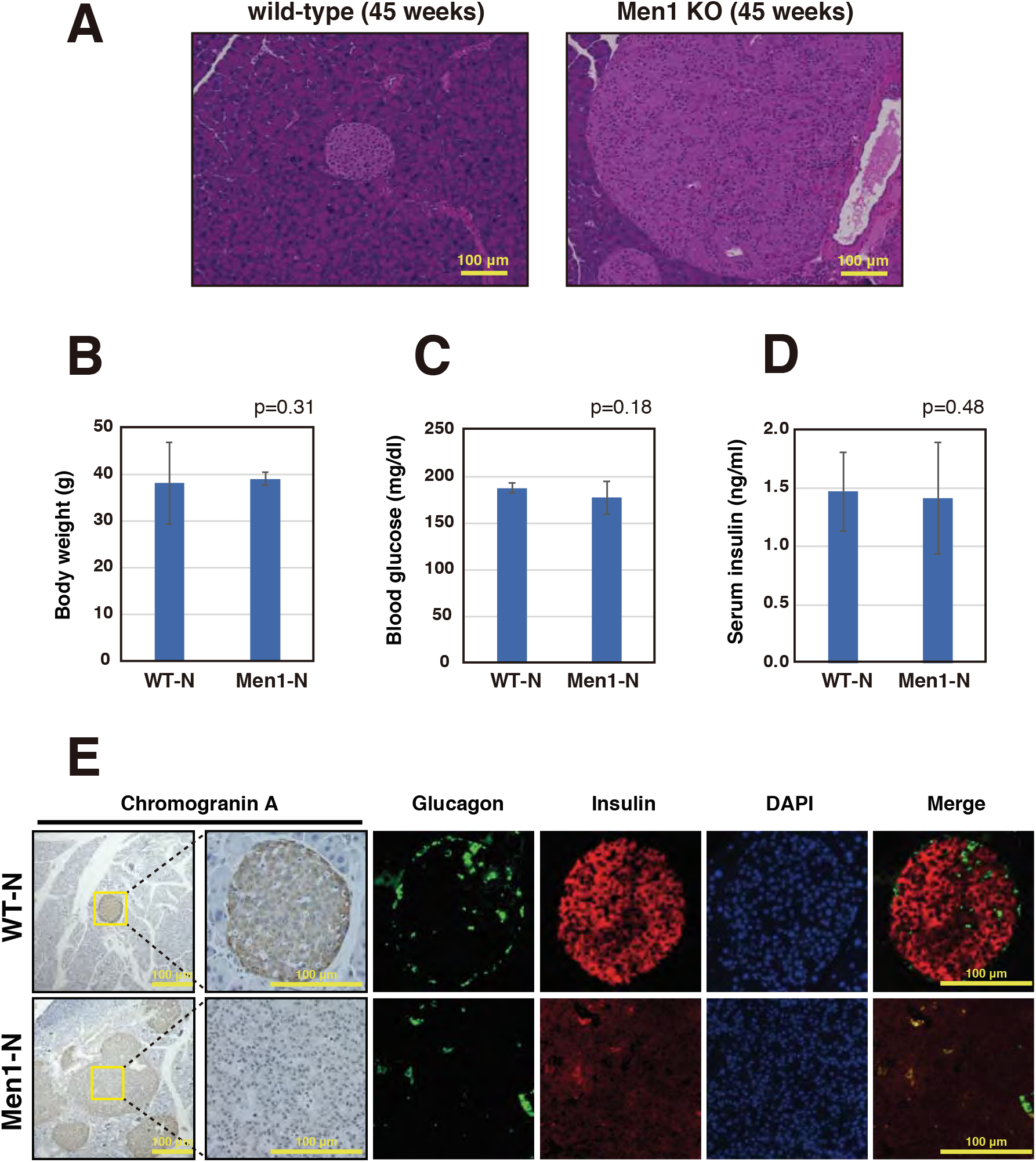
β cell-specific *Men1* knock-out mice develop non-functional PanNETs. **A.** Pancreatic sections of 45-week-old wild-type mice and *Men1*^f/f^-RipCre^+^ mice fed a normal diet were stained with HE. **B, C.** Body weights (B) and blood glucose levels (C) of 45-week-old wild-type (N=19) or *Men1*^f/f^-RipCre^+^ (N=21) mice fed a normal diet. Mice were fed ad libitum. Data are represented as the mean fold expression ± standard error of the mean (SEM). **D.** Serum insulin levels of 45-week-old wild-type (N=19) or *Men1*^f/f^-RipCre^+^ mice fed a normal diet (N=18). Mice were fed ad libitum. Data are represented as the mean fold expression ± SEM. **E.** Pancreatic sections of 45-week-old wild-type or *Men1*^f/f^-RipCre^+^ mice fed a normal diet were stained with anti-insulin, anti-glucagon and anti-Chromogranin A.

### A ketogenic diet suppresses PanNET development

We found that male mice lacking *Men1* exons 3-6 in β cells develop PanNET by the age of 45 weeks. These mice have almost no abnormalities at 10 weeks (Figs. 2A-2C). In order to analyze the effect of a ketogenic diet on non-functional PanNET development, we started feeding the *Men1*^f/f^-RipCre^+^ male mice with a ketogenic diet from the age of 7-13 weeks. The mice were sacrificed at age 45 weeks and the sizes of the islets were analyzed. As shown in Figs. 2A-2C and S1, the ketogenic diet significantly suppressed the development of islet hyperplasia and PanNET. There was no significant difference in the sizes of the islets of the ketogenic diet-fed mice and wild-type mice at the same age (Fig. 2C). In addition, the sizes of the islets of *Men1*^f/f^-RipCre^+^ mice fed the ketogenic diet were not significantly different from those of 10-week-old mice, showing that the ketogenic diet suppressed the enlargement of the islets during the first 35 weeks (Fig. 2C).

**Fig. 2.**
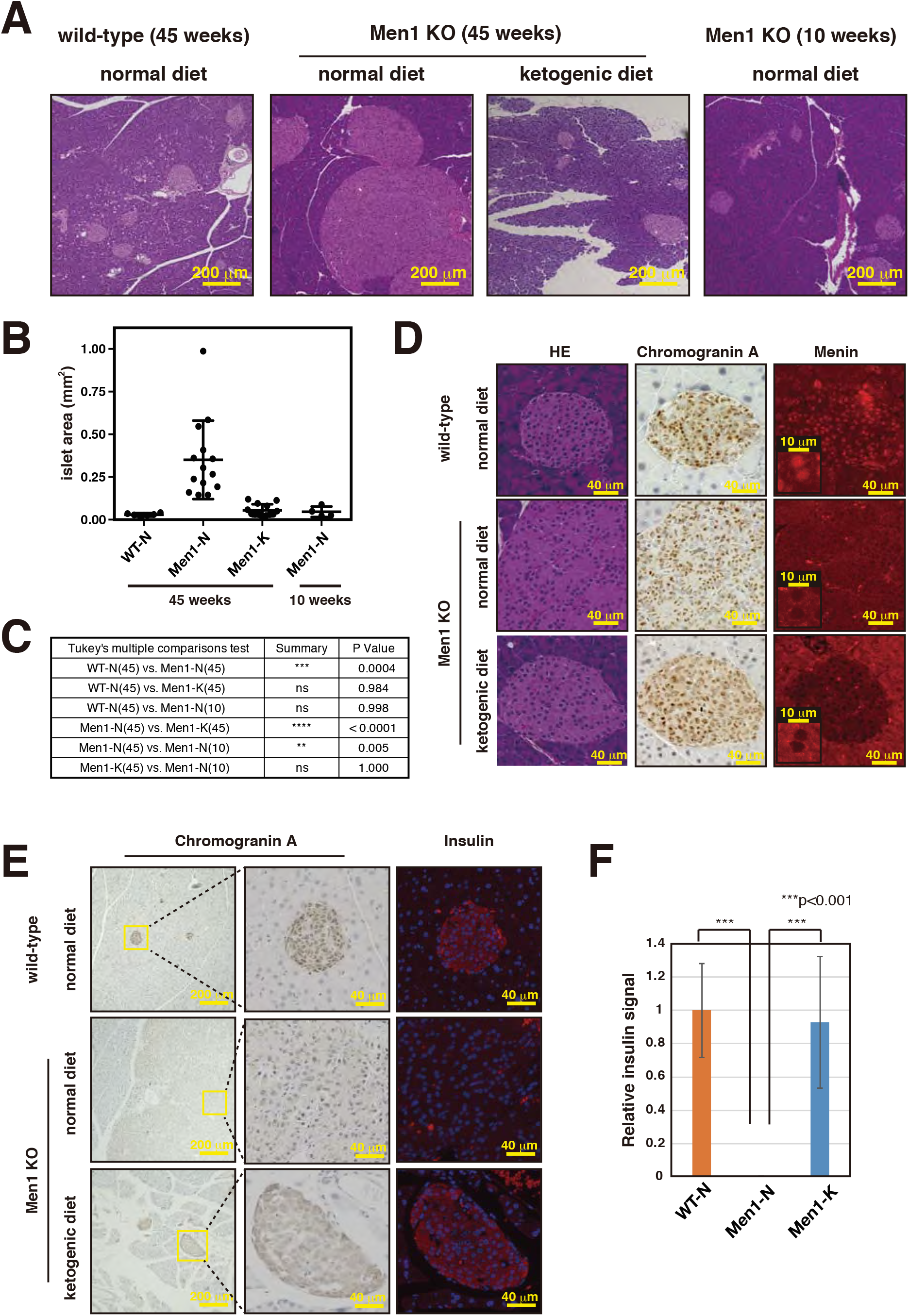
Ketogenic diet suppresses non-functional PanNET development. **A.** Pancreatic sections of 45-week-old wild-type mice fed a normal diet, *Men1*^f/f^-RipCre^+^ mice fed a normal diet or *Men1*^f/f^-RipCre^+^ mice fed a ketogenic diet from 8 weeks of age were stained with HE. HE staining of pancreatic sections of 10-week-old *Men1*^f/f^-RipCre^+^ mice fed a normal diet is also shown. **B.** Islet areas of 45-week-old wild-type mice fed a normal diet (N=6), *Men1*^f/f^-RipCre^+^ mice fed a normal diet (N=14) or *Men1*^f/f^-RipCre^+^ mice fed a ketogenic diet from 7-13 weeks of age (N=13), and 10-week-old *Men1*^f/f^-RipCre^+^ mice fed a normal diet (N=4) were analyzed. The areas of the top 10 largest islets were calculated and the means of the 10 islet areas were obtained for each mouse. Data are represented as the mean fold expression ± standard deviation (SD). **C.** Statistical analysis of the average islet sizes of each group of mice was performed. P values were obtained using the one-way ANOVA test. The age of the mice sacrificed are shown in parenthesis. **D.** Pancreatic sections of 45-week-old wild-type mice fed a normal diet, *Men1*^f/f^-RipCre^+^ mice fed a normal diet or ketogenic diet from 8 weeks of age were stained with HE, anti-Chromogranin A and anti-Menin. **E, F.** Pancreatic sections of 45-week-old wild-type mice fed a normal diet, *Men1*^f/f^-RipCre^+^ mice fed a normal diet or ketogenic diet from 8 weeks of age were stained with anti-insulin and anti-Chromogranin A. Relative intensities of insulin staining from 3 mice of each group (total 9-12 islets) were calculated and shown in Graph F. Data are represented as the mean fold expression SD.

We next analyzed the features of the islets of the ketogenic diet-fed *Men1*^f/f^-RipCre^+^ mice. As shown in Fig. 2D, islets from *Men1*^f/f^-RipCre^+^ mice fed either a normal or ketogenic diet were both positive for the neuroendocrine marker chromogranin A and had lost the expression of Menin as expected. As shown in Fig. 1E, *Men1*-deficient islets at 45 weeks of age had decreased insulin expression. However, this decrease in islet insulin expression was rescued in mice fed a ketogenic diet (Figs. 2E and 2F). Collectively these results show that the ketogenic diet dramatically suppresses the onset of non-functional PanNET.

### A ketogenic diet suppresses pancreatic islet cell proliferation

The results indicate that the lower blood glucose levels resulting from a ketogenic diet also resulted in lower insulin levels in the blood and decreased insulin signaling in islet cells. We first analyzed the body weights and blood glucose levels in 45-week-old mice fed a normal versus ketogenic diet ad libitum. As shown in Figs. 3A and 3B, blood glucose levels were significantly lower in the mice fed a ketogenic diet, while there were no significant differences in body weights. We next performed a food tolerance test using fasted mice and analyzed blood glucose and serum insulin levels 1.5 hrs post food intake. In the case of wild-type mice, blood glucose levels continue to decrease during the first 48 hours of fasting, while liver glycogen is not completely depleted until 36 hours of fasting ^26, 27^. Based on these findings, we implemented a fasting period from overnight to 41 hours in our study to minimize the effects of endogenous glucose. While there was a significant increase in blood glucose and high insulin secretion in the mice refed a normal diet, these increases in blood glucose and insulin were not detected in mice fed a ketogenic diet (Figs. 3C, 3D and S2). It was also shown that the insulin-regulated PI3K-Akt-mTOR pathway (P-S6 and P-mTOR) is attenuated in the islet cells of mice fed a ketogenic diet (Figs. 3E-3H). Since insulin signaling is closely related to islet cell proliferation, it is expected that decreased insulin levels in the blood should result in decreased islet cell proliferation. As shown in Figs. 3I-3K, the number of Ki67-positive proliferating cells is lower and the expression of p27, a representative cell cycle inhibitor, was higher in the islets of mice fed a ketogenic diet. The *Cdkn1b* gene encoding p27 is a target gene of Menin, and its mRNA expression was not affected by the ketogenic diet, suggesting that p27 expression is regulated at the post-transcriptional level (Fig. S3). These data show that the ketogenic diet suppressed the proliferation of *Men1*-deficient islet cells.

**Fig. 3.**
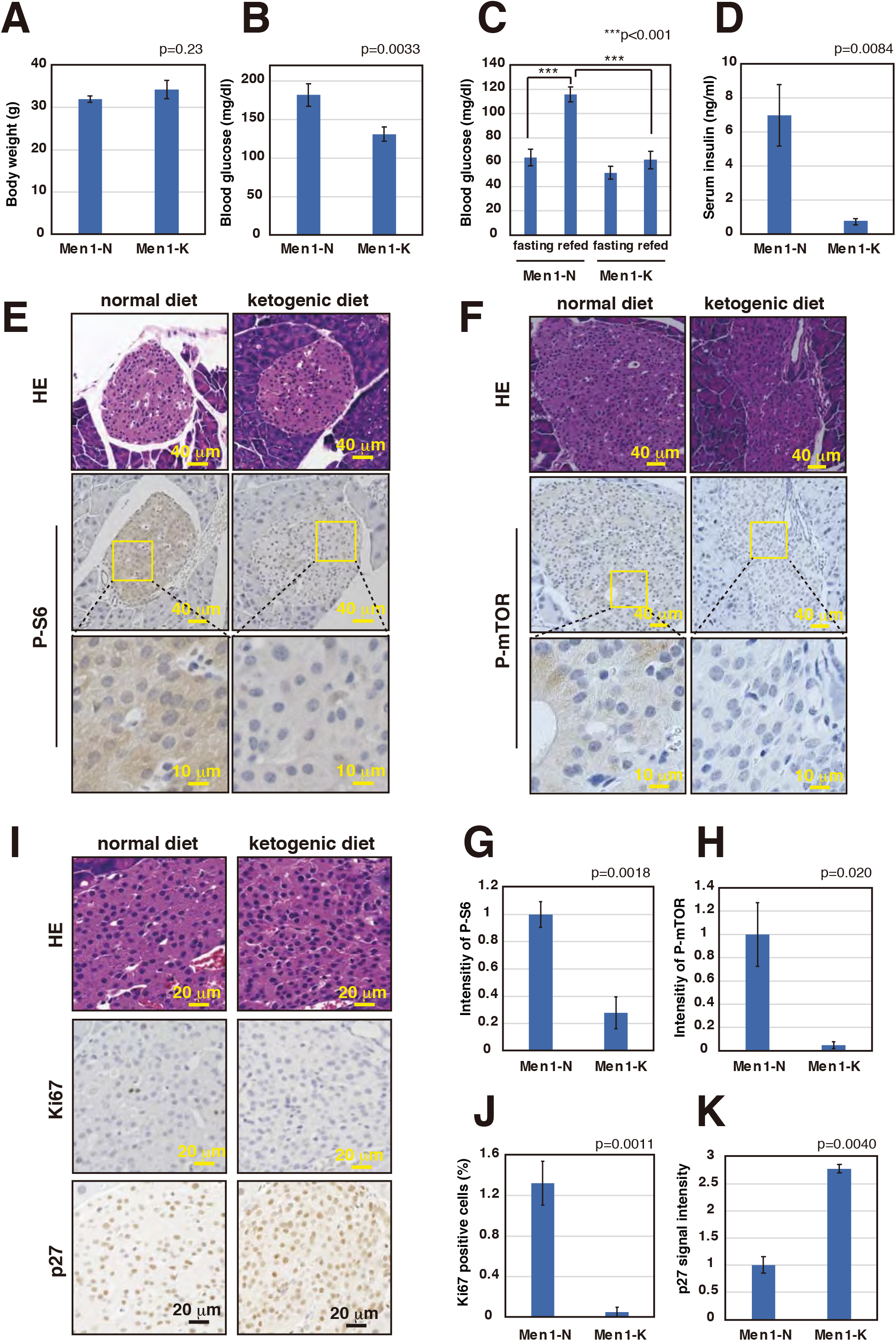
Ketogenic diet suppresses pancreatic islet cell proliferation. **A.** Body weights (A) and blood glucose levels (B) of 45-week-old *Men1*^f/f^-RipCre^+^ mice fed a normal diet (N=18) or ketogenic diet (N=32). Mice were fed ad libitum. Data are represented as the mean fold expression ± SEM. **C, D.** Blood glucose levels (C) and serum insulin levels (D) of 20-week-old *Men1*^f/f^-RipCre^+^ mice fed a normal diet (N=4) or ketogenic diet from 10 weeks of age (N=3). Fasted mice (mice fasted for overnight to 41 hrs) and mice refed the usual meal for 1.5 hrs were analyzed. Data are represented as the mean fold expression ± SEM. **E-G.** Mice were treated as in C and pancreatic sections of mice refed a normal diet or ketogenic diet were stained with HE, anti-P-S6 (E) and anti-P-mTOR (F). High magnification images are shown in the bottom panels. Relative intensities of P-S6 staining from 5 (normal diet) and 4 (ketogenic diet) islets were calculated and shown in graph G. Relative intensities of P-mTOR staining from 4 (normal diet) and 4 (ketogenic diet) islets were calculated and shown in graph H. Data are represented as the mean fold expression ± SEM. **I-K.** Pancreatic sections of 30-week-old *Men1*^f/f^-RipCre^+^ mice fed a normal diet or ketogenic diet from 10 weeks of age were stained with HE, anti-Ki67 and anti-p27 (I). Total and Ki67 positive nuclei numbers from 4 islets were counted and shown in Graph J. Relative intensities of p27 staining from 3 islets were calculated and shown in Graph K. Data are represented as the mean fold expression ± SEM.

### A ketogenic diet changes microenvironment of the islet

It has been reported that angiogenesis is an important step in PanNET tumorigenesis and angiogenic switch is turned on upon progression from hyperplasia to neoplasia ^28^. Insulin signaling enhances angiogenesis in the islets and PI3K-Akt-mTOR pathway has been shown to regulate angiogenesis ^29 30^. Since we found that PI3K-Akt-mTOR pathway activation is suppressed by a ketogenic diet, we analyzed whether a ketogenic diet reverses the vascularized phenotype of the PanNETs formed in *Men1*^f/f^-RipCre^+^ mice. As shown in Fig. 4A and 4B, while large blood vessels were frequently found in the islets of the mice fed a normal diet, such vessels were highly reduced in the islets of the mice fed a ketogenic diet. We further found that expression of angiogenic genes (Mmp-2 and Mmp-9) were down regulated in the islets of the mice fed a ketogenic diet (Figs. 4C and 4D). Since it is expected that suppression of angiogenesis in the islets will further change the microenvironment of the islets, we further analyzed the expression of genes involved in macrophage activation (Il-1β and Nos2). Expression of both genes were down regulated by ketogenic diet showing that macrophage activation was suppressed by the diet (Figs. 4E and 4F). On the other hand, expression of the endocrine marker Synaptophysin was not significantly different among the samples, indicating consistency in islet isolation (Fig. 4G). Collectively, data show that a ketogenic diet suppresses angiogenesis and macrophage activation, two important microenvironment involved in PanNET progression.

**Fig. 4.**
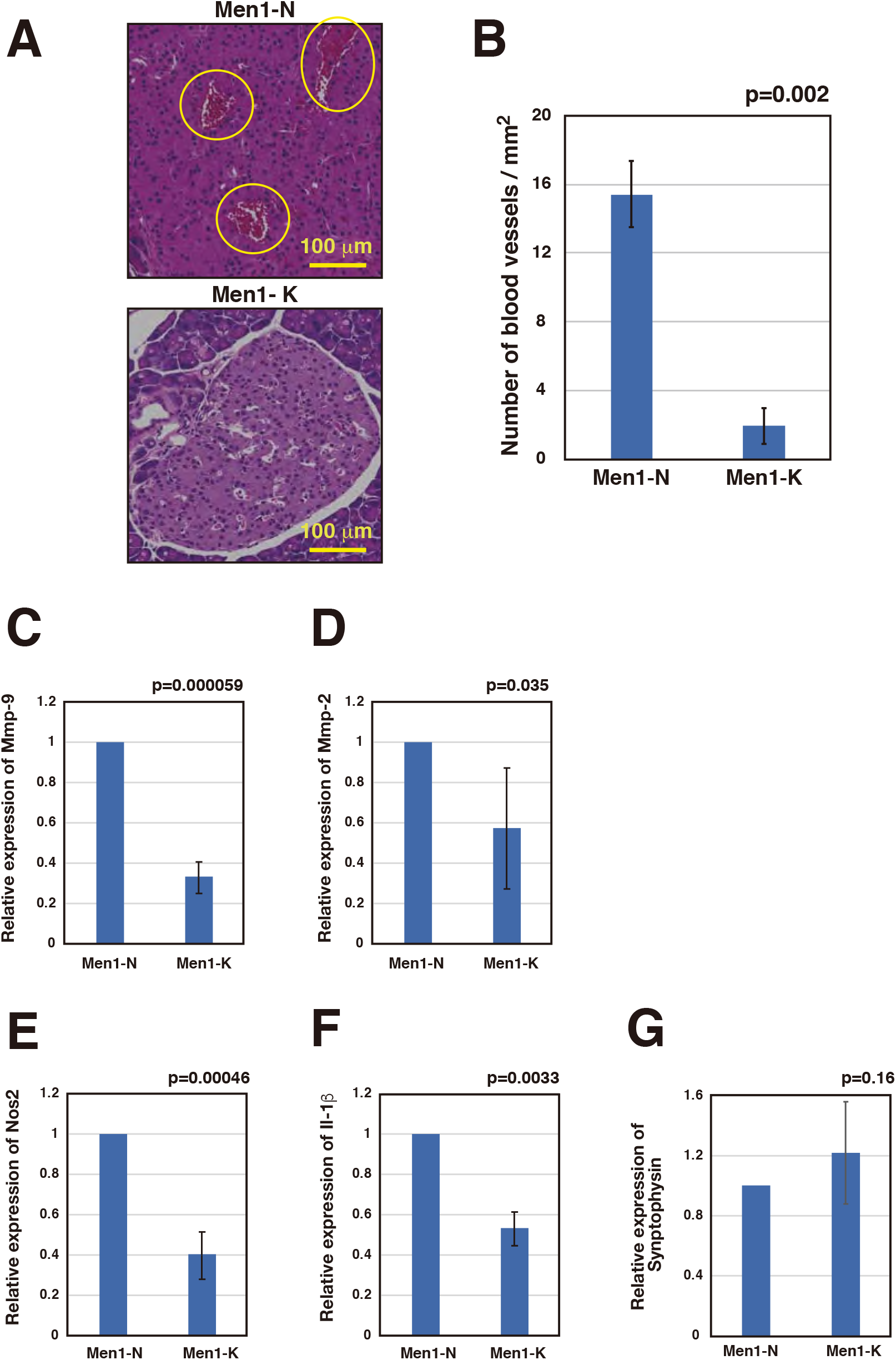
Ketogenic diet suppresses angiogenesis and macrophage activation. **A.** HE stained pancreatic sections of 45-week-old *Men1*^f/f^-RipCre^+^ mice fed a normal diet (N=3) or *Men1*^f/f^-RipCre^+^ mice fed a ketogenic diet from 11 weeks of age (N=3) were analyzed. Typical vascularized islet from the mice fed a normal diet and a non-vascularized islet from the mice fed a ketogenic diet are shown (A). The number of vessels with an area of 0.0006 mm^2^ or greater was calculated from 4 fields derived from 4 islets from each mouse. The numbers of vessels are counted and shown as a graph in B. Data are represented as the mean fold expression ±SD. **C-G.** Islets were isolated from 45-week-old *Men1*^f/f^-RipCre^+^ mice fed a normal diet or *Men1*^f/f^-RipCre^+^ mice fed a ketogenic diet from 10 weeks of age. 3 sets of islets were isolated from the mice fed a normal diet (N=2, 2, 3) or mice fed a ketogenic diet from 10 weeks of age (N=2, 3, 7). mRNAs were purified from islets and the expression of genes involved in angiogenesis (*Mmp2* and *Mmp9*) (C, D) and macrophage activation (*Nos2* and *Il1β*) (E, F) were analyzed. An endocrine marker, Synaptophysin, was also analyzed to evaluate the consistency of islet isolation (G). Data are represented as the mean fold expression ± SD.

### A ketogenic diet suppresses PanNET progression

As the ketogenic diet was found to suppress the onset and development of PanNETs in *Men1*^f/f^-RipCre^+^ mice, we next analyzed whether this diet also suppresses the progression of PanNETs. At 30 weeks of age, *Men1*-deficient islets were found to be significantly larger than wild-type islets (Fig. 5A): the average size of the largest 10 islets of *Men1*^f/f^-RipCre^+^ mice at 30 weeks of age was 0.17 mm^2^, while those from wild type mice averaged 0.028 mm^2^. We therefore started feeding the 30-week-old *Men1*^f/f^-RipCre^+^ mice a ketogenic diet. When the islet sizes were subsequently analyzed at age 45 weeks, the islet sizes of the mice fed a ketogenic from 30 weeks were significantly smaller than those of mice fed a normal diet (Figs. 5B-5D and S4). Furthermore, the islet sizes were not significantly different from those of 30-week-old *Men1*^f/f^-RipCre^+^ mice, showing that the ketogenic diet suppressed the progression of PanNET for 15 weeks when the mice were fed this diet. We also analyzed the effect of the diet on islet cell proliferation and angiogenesis. As shown in Figs. 5E-5H, the ketogenic diet significantly suppressed islet cell proliferation and angiogenesis further, indicating that the diet significantly suppressed PanNET progression.

**Fig. 5.**
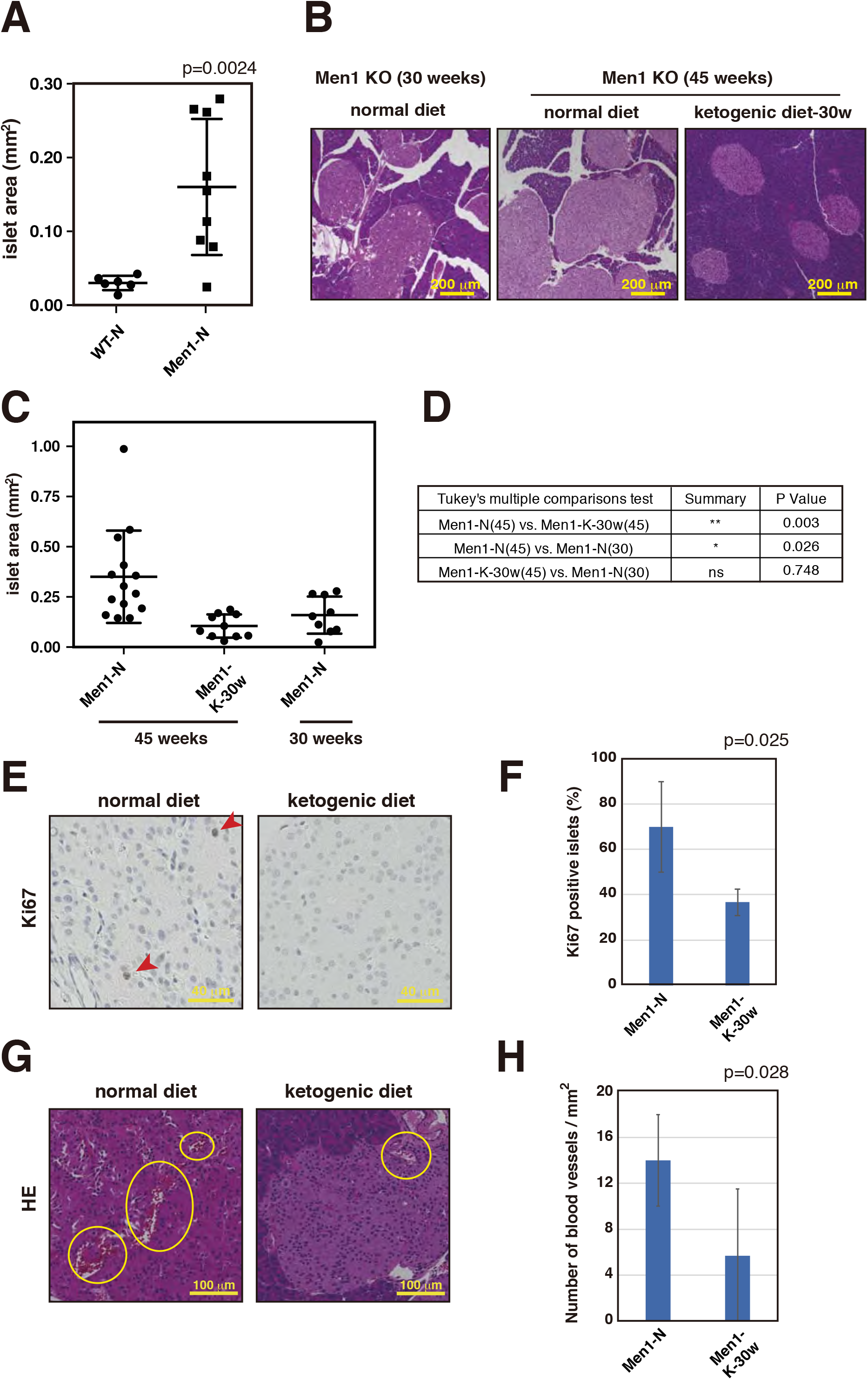
Ketogenic diet suppresses PanNET progression. **A.** Islet areas of 30-week-old wild-type mice and *Men1*^f/f^-RipCre^+^ mice fed a normal diet were analyzed. The areas of the top 10 largest islets were calculated and the means of the 10 islet areas were obtained for each mouse. Data are represented as the mean fold expression ± SD. **B.** Pancreatic sections of 45-week-old *Men1*^f/f^-RipCre^+^ mice fed a normal diet or ketogenic diet from 30 weeks of age were stained with HE. HE staining of pancreatic sections of 30-week-old *Men1*^f/f^-RipCre^+^ mice fed a normal diet is also shown. **C.** Islet areas of 45-week-old *Men1*^f/f^-RipCre^+^ mice fed a normal diet or ketogenic diet from 30 weeks of age, and 10- or 30-week-old *Men1*^f/f^-RipCre^+^ mice fed a normal diet were analyzed as in A. *Men1*^f/f^-RipCre^+^ mice fed a normal diet is same as the mice described in Fig. 5A. Data are represented as the mean fold expression ± SD. **D.** Statistical analysis of the average islet sizes of each group of mice was performed. P values were obtained using the one-way ANOVA test. The age of the mice sacrificed are shown in parenthesis. **E, F.** Pancreatic sections of 45-week-old *Men1*^f/f^-RipCre^+^ mice fed a normal diet (N=3) or ketogenic diet from 30 weeks of age (N=3) were stained with anti-Ki67 (E). Graph F shows the number of total versus Ki67-positive nuclei from 4 islets. Data are represented as the mean fold expression ±SD. **G, H.** HE-stained pancreatic sections of 45-week-old *Men1*^f/f^-RipCre^+^ mice fed a normal diet (N=4) or *Men1*^f/f^-RipCre^+^ mice fed a ketogenic diet from 30 weeks of age (N=4) were analyzed as shown in Fig. 4B. The numbers of vessels are shown in graph H. Data are represented as the mean fold expression ±SD.

### A ketogenic diet suppresses pituitary NET development

*Men1*^f/f^-RipCre^+^ mice develop not only PanNETs but also pituitary NETs. Pituitary NET development is more enhanced in female mice and most of the female mice develop pituitary NETs by the age of 45 weeks. We therefore analyzed the effect of a ketogenic diet on pituitary NET development in 45-week-old female *Men1*^f/f^-RipCre^+^ mice. We started feeding the *Men1*^f/f^-RipCre^+^ female mice with a ketogenic diet from the age of 10-13 weeks. As shown in Figs. 6A-6C, a ketogenic diet significantly suppressed pituitary NET development in these mice. It has been reported that *Men1*-deficient mice frequently develop prolactin-positive pituitary NETs. This previous report was confirmed, and most of the *Men1*-deficient mice fed a normal diet developed prolactin-positive tumors (Figs. 6D and 6E). Within the pituitary NET of the mouse fed a normal diet, accumulation of prolactin-positive cells was observed, while no cells were positive for TSH, another pituitary hormone. On the other hand, the distribution of prolactin- and TSH-producing cells were similar to those of wild type mice, and the clonal expansion of prolactin- and TSH-producing cells was not evident in the pituitaries of mice fed a ketogenic diet from 10 weeks of age (Figs. 6D and 6E).

**Fig. 6.**
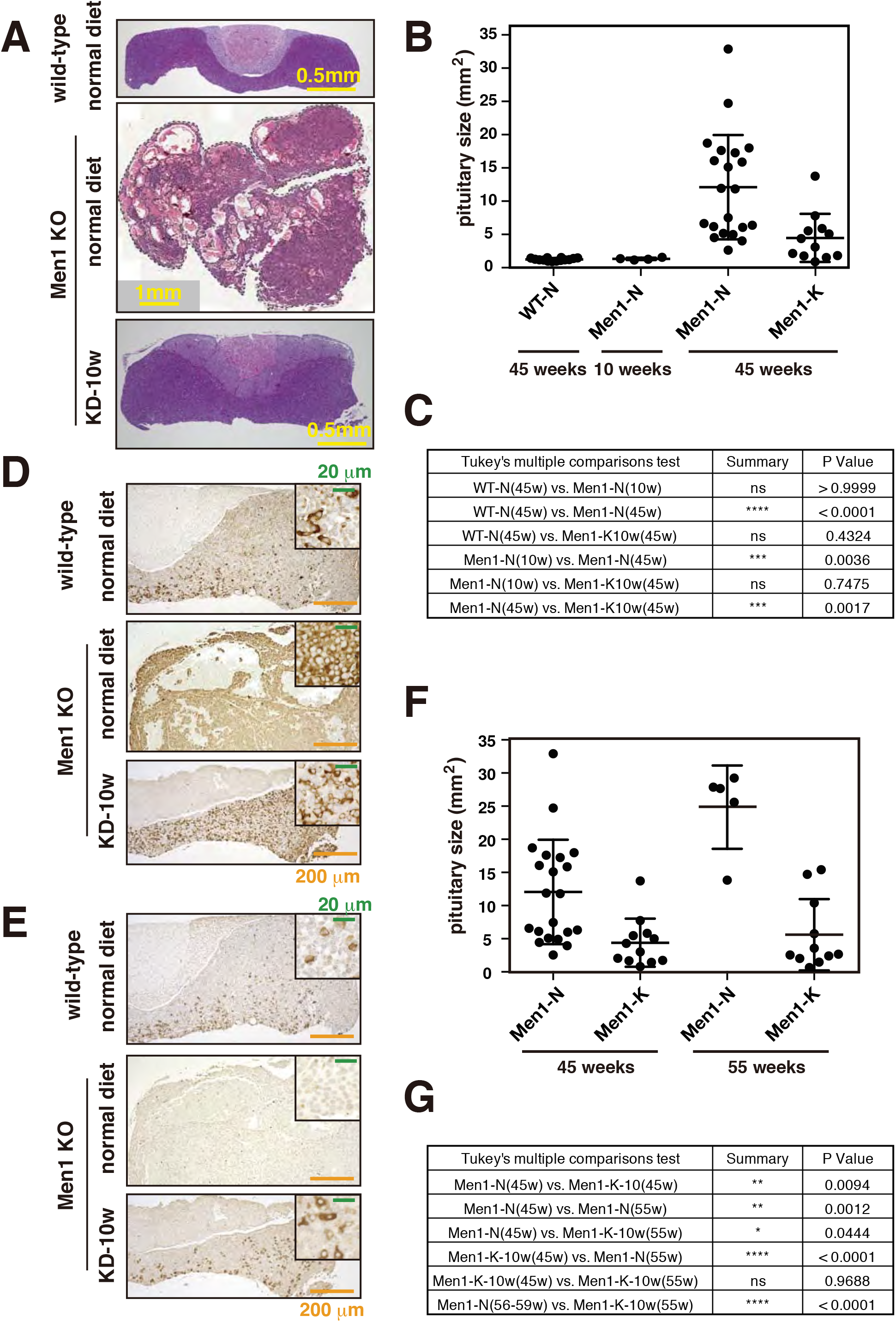
Ketogenic diet suppresses pituitary NET development. **A.** Pituitary sections of 45-week-old wild-type mice fed a normal diet, *Men1*^f/f^-RipCre^+^ mice fed a normal diet or ketogenic diet from 10 weeks of age were stained with HE. **B.** Pituitary sizes of 45-week-old wild-type mice fed a normal diet, *Men1*^f/f^-RipCre^+^ mice fed a normal diet or ketogenic diet from 10-13 weeks of age were analyzed. Pituitaries of 10-week-old *Men1*^f/f^-RipCre^+^ mice fed a normal diet were also analyzed. Data are represented as the mean fold expression ± SD. **C.** Statistical analysis of the pituitary sizes of each group of mice was performed. P values were obtained using the one-way ANOVA test. The age of the mice sacrificed are shown in parenthesis. **D, E.** Pituitary sections of 45-week-old wild-type mice fed a normal diet and *Men1*^f/f^-RipCre^+^ mice fed a ketogenic diet from 13 weeks of age were stained with anti-prolactin (D) and anti-TSH (E) antibodies. **F.** Pituitary sizes of 55-week-old *Men1*^f/f^-RipCre^+^ mice fed a normal diet or ketogenic diet from 10-13 weeks of age were analyzed and compared to those of 45-week-old mice. The data for the 45-week-old mice is the same as in Fig. 6B. Data are represented as the mean fold expression ± SD. **G.** Statistical analysis of the pituitary sizes of each group of mice was performed as in C.

As shown in Fig. S5, the median survival of female *Men1*^f/f^-RipCre^+^ mice is 56 weeks, and these mice die from pituitary NETs. We, therefore, fed the mice a ketogenic diet to the age of 55 weeks and analyzed the size of the pituitaries. As shown in Figs. 6F and 6G, ketogenic diet significantly suppressed the pituitary NET development until the age of 55 weeks. The sizes of the pituitaries of 55-week-old mice were similar to those of the 45-week-old mice. Collectively, these results show that a ketogenic diet also suppressed pituitary NET development and contribute to the extension of the life of these mice.

### The relationship between PanNET patient prognosis and blood glucose level

We next analyzed the relationship between overall survival and blood glucose levels in 54 patients who had been pathologically diagnosed as having non-functional PanNETs (Grade 1-3) and those who had undergone chemotherapy. Patients who developed multiple primary tumors were excluded from the analysis. The American Diabetes Association has determined that a fasting blood glucose level from 100 to 125 mg/dl is an indication of a prediabetic state ^31^. Therefore, high blood glucose levels were defined in our study as blood glucose levels higher than 100 mg/dl. The median blood glucose levels in the low and high blood glucose groups were 92 mg/dl (76–100 mg/dl) and 117 mg/dl (101– 245 mg/dl), respectively, and in the high blood glucose level group, 16 (48%) of 33 patients had blood glucose level greater than 125 mg/dl. As shown in Fig. 7A, there was a tendency for patients with lower blood glucose levels to have a better prognosis (p=0.080). On the other hand, when major clinicopathological characteristics are classified into low and high blood glucose level groups, as listed in Fig. 7B, the distributions of age, sex and tumor grade were not significantly different between the two groups. For patients who had undergone chemotherapy, there was no significant bias between the blood glucose high and low groups for the various chemotherapeutic regimens, also listed in Fig. 7C. Collectively, these data suggest that high blood glucose induces the progression of PanNETs.

**Fig. 7.**
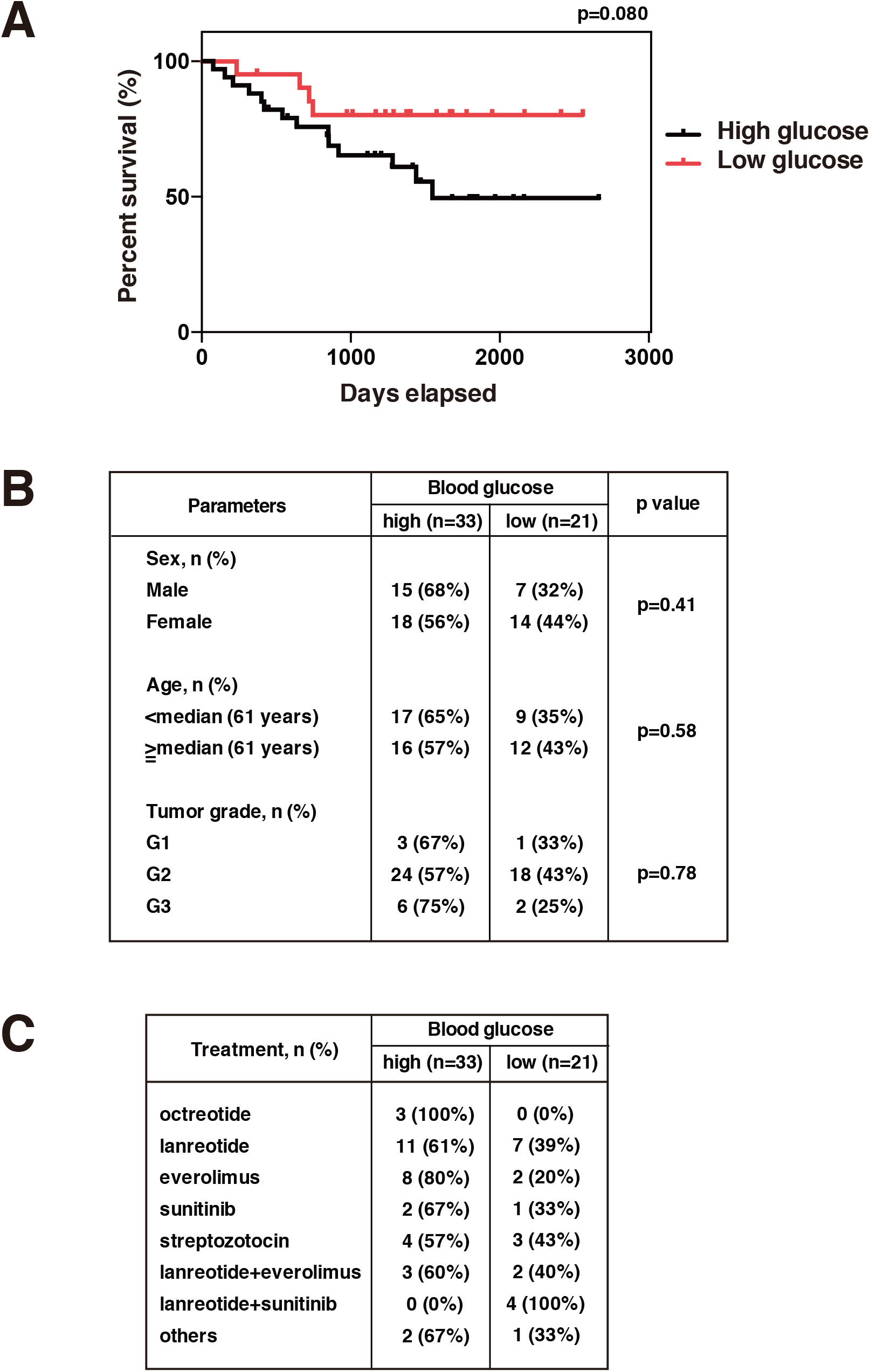
The prognosis of PanNET patients having lower blood glucose levels are better than those having higher blood glucose levels. **A.** Overall survival of patients with high and low fasting blood glucose levels (N=54). **B.** Baseline characteristics of patients analyzed are shown. **C.** Chemotherapeutic regimens of patients analyzed are shown.

## Discussion

In this study, we have shown that the metabolic intervention of a very low carbohydrate diet (ketogenic diet) could significantly suppress both the initiation and progression of non-functional PanNET and pituitary NET in a mouse model. Recent advances in the understanding of the molecular factors involved in pancreatic and pituitary NETs have led to new antitumor treatments, including reagents acting on specific molecular targets. For example, inhibitors of receptor tyrosine kinases and mTOR are in clinical use for the treatment of PanNETs, however they may have limited efficacy and patients often stop responding to these inhibitors ^32^. The most frequently used PanNET mouse model used for preclinical testing of therapies is a RipTag2 mouse model, which expresses the SV40 T antigen under the control of insulin promoter and develops functional PanNETs ^33^. There are also several independent β cell-specific *Men1* knock-out mice that develop functional insulinomas ^7, 8 9^. With these functional PanNET mouse models, it has been shown that several molecular targeted drugs such as an anti-VEGF monoclonal antibody (targeting the angiogenic pathways) or nintedanib (targeting VEGF, PDGF, and fibroblast growth factor) have anti-tumorigenic effects ^34 35 36^. However, there have been no preclinical studies involving mouse models of non-functional PanNETs. We have established a non-functional PanNET mouse model and shown that by maintaining low insulin signaling by a ketogenic diet, it is possible to prevent and suppress the progression of non-functional PanNETs. We have shown that this dietary intervention is also effective for pituitary NETs. Recently, it has been reported that the ketogenic diet may elicit its anticancer effect by acting on the cancer epigenome ^37^. Analysis of ketogenic-diet-induced changes in the cancer epigenome is an interesting issue to be analyzed in future studies. It has also been reported that a low-carbohydrate ketogenic diet is safe and has no adverse effects in mice and can improve the effectiveness of some anticancer therapies ^15 16 17^. Therefore, it would be of great interest in future studies to test combinations of a ketogenic diet with other anticancer therapies used for PanNETs and pituitary NETs.

The results shown in this study can be translated not only to sporadic but also to familial non-functional PanNETs and pituitary NETs. In MEN1 patients that develop familial NETs, non-functional PanNETs are highly prevalent and are the major cause of premature MEN1-related death^32^. Surgery has been the only curative option for nonfunctioning PanNETs, however, since MEN1 patients often have multiple PanNETs that occur at a younger age and have a higher metastatic potential, surgery is not always possible. Although clinical guidelines recommend early detection by genetic testing of asymptomatic relatives of MEN1-mutated patients within the first decade of life, there is currently no effective method of prevention for these patients. However, this preclinical study shows that dietary intervention with a very low carbohydrate diet is effective in preventing nonfunctioning PanNET in mice, and this method may be applicable to MEN1 patients.

This dietary intervention is effective not only for the prevention of PanNETs but also for the suppression of PanNET progression, and has no adverse side effects. We also showed that higher blood glucose levels are related to poorer prognosis of PanNET patients that are receiving chemotherapy. It has also been reported that high pre-operative blood glucose predicts a poor prognosis in PanNET patients ^38^. In addition, metformin, a drug that reduces blood glucose levels, enhances the effect of chemotherapy in PanNET patients ^39^. These results suggests that by lowering blood glucose, PanNET progression can also be suppressed in humans. Future clinical studies to investigate the effectiveness of this method for both familial and sporadic human non-functional PanNETs and pituitary NETs should be of great interest. It is also possible that this treatment could be applied to other types of cancers including various familial cancers that are dependent on the insulin-regulated PI3K-Akt-mTOR pathway, and could represent the first effective prevention method that does not have adverse effects.

## Materials and Methods

### Mice used in this study

Mouse experiments were performed in a specific pathogen-free environment at the National Cancer Center animal facility according to institutional guidelines and all the animal experiments were approved by the Committee for Ethics in Animal Experimentation at the National Cancer Center (T17-002-C01 and T17-011-M03). Mice were kept in an animal facility controlled at a temperature of 22±0.5°C, 45〜65 % humidity, pressure difference of +10 to 20 Pa, ventilation of 15 or more air volumes per hour, class 10,000 clean room, 150 to 300 lux illumination (40 to 85 cm above the breeding room floor) and a 12 h light and dark cycle (8 am to 8 pm; light). Pancreatic β-cell-specific *Men1* deficient mice were generated by crossing mice carrying the floxed allele (*Men* ^f/f^) with RIP-Cre transgenic mice expressing the Cre recombinase under the control of the rat insulin II promoter (B6.Cg-Tg(Ins2-cre)25Mgn/J), termed *Men1*^f/f^-RipCre^+^ mice ^25 40^. The mice were backcrossed against C57BL6 at least 8 times before use in experiments. *Men1*^f/f^-RipCre^+^ mice develop both PanNETs and pituitary NETs. While PanNET development is enhanced in male mice, pituitary NET development is more enhanced in female mice. By 45 weeks of age, most of the male mice develop PanNETs and most of the female mice develop pituitary NETs. We therefore analyzed the effect of a ketogenic diet on PanNETs in male mice and pituitary NETs in female mice. Mice were fed a normal diet (CE-2, Crea Japan, Tokyo, Japan) or Ketogenic AIN-76A-Modified Diet, High Fat, Paste (Bio-serv, NJ).

### Blood glucose and Plasma Insulin Measurement

Blood measurements were performed at 1-2pm. Blood glucose levels were determined with blood samples from tail vein punctures in mice using a Glucose pilot instrument (Aventir Biotech, Carlsbad, CA), according to the manufacturer’s protocol. Plasma insulin levels were determined with blood samples from the inferior vena cava in mice using the Morinaga Ultra Sensitive Mouse Insulin Elisa Kit (Morinaga Institute of Biological Science, Inc., Tokyo, Japan), according to the manufacturer’s protocol.

### Immunohistochemistry (IHC)

IHC was performed basically as described according to the manufacturer’s instructions ^20^. In brief, after deparaffinization, tissues sections underwent antigen retrieval by autoclaving slides in 10 mM citrate buffer (pH 6.0). Sections were pretreated with 3 % H_2_O_2_ for inactivation of endogenous peroxidase. The primary antibodies used in the study were; rabbit anti-Ki67 monoclonal antibody (SP6, Novus Biologicals) diluted 1:200, rabbit anti-Chromogranin A polyclonal antibody (Thermo Scientific) diluted 1:200, rabbit anti-P-S6 (#5364, CST) diluted 1:500, rabbit anti-P-mTOR (#2976, CST) diluted 1:100, rabbit anti-p27 (C19, Santacruz) diluted 1:1000, rabbit anti-rat Prolactin (the Institute for Molecular and Cellular Regulation, Gunma University) diluted 1:5000, rabbit anti-rat TSH (the Institute for Molecular and Cellular Regulation, Gunma University) diluted 1:5000. As secondary antibodies, biotinylated anti-rabbit IgG antibody (VECTOR Laboratories) were used. We also used SignalStain Boost Detection Reagent (HRP, Rabbit #8114) for CST antibodies. We used 3,3’-diaminobenzidine tetrahydrochloride (DAB; Muto Pure Chemicals) as the substrate chromogen. The sections were counter-stained with hematoxylin. For fluorescent immunohistochemical staining of insulin, glucagon and Menin, nonspecific interactions were blocked for 1h using a goat serum solution. The primary antibodies were: guinea pig anti-insulin polyclonal antibody (Abcam) diluted 1:400, mouse anti-glucagon monoclonal antibody (SIGMA-ALDRICH) diluted 1:750, rabbit anti-Menin polyclonal antibody (Bethyl Laboratories, A300-105A) diluted 1:3000 with Signal Enhancer HIKARI (NACALAI TESQUE). These were applied to the slides and incubated overnight at 4 °C. As secondary antibodies, Alexa Fluor 546 goat anti-rabbit IgG antibody (Invitrogen) diluted 1:500 and Alexa Fluor 546 or 594 goat anti-guinea pig IgG antibody (Invitrogen) diluted 1:1000 with PBST-BSA were applied to the slides and incubated 3 hrs at RT. For mouse antibodies, the Vector MOM immunodetection kit was used based on the protocol specified by the manufacturer (Vector Laboratories).

### Isolation of Mouse Islets

Isolation of mouse islets was performed as described ^20^. Briefly, mouse islets were isolated from 45-week-old male animals by collagenase digestion of the pancreas, followed by purification using a Ficoll gradient. Islets were handpicked and used for the subsequent analysis as shown in Figs. 4C-4G and S3.

### Reverse Transcription and Real-Time PCR

Total cellular RNA was extracted using AllPrep DNA/RNA/Protein mini kit (Qiagen, UK). Reverse transcription was carried out using kits from Invitrogen following the manufacturer’s instructions (SuperScript First-Strand Synthesis System for RT-PCR). Total RNA (0.2–5 μg) was used for reverse transcription. Reverse-transcribed cDNAs were subjected to realtime PCR, which was performed for genes of interest by using the SYBR Green Premix ExTaq (Tli RNaseH Plus) (Takara bio Inc, Japan) and the CFX96 Real-time PCR (Biorad, UK), according to manufacturer’s instructions. TaqMan probe for mouse 18S rRNA, β-actin and Il-1β from Integrated DNA Technologies and custom-designed primers were used to detect Cdkn1b (F; AGCAGTGTCCAGGGATGAGGAA, R; TTCTTGGGCGTCTGCTCCACAG), Mmp2 (F; CAAGGATGGACTCCTGGCACAT, R; TACTCGCCATCAGCGTTCCCAT), Mmp9 (F; GCTGACTACGATAAGGACGGCA, R; TAGTGGTGCAGGCAGAGTAGGA), Nos2 (F; GAGACAGGGAAGTCTGAAGCAC, R; CCAGCAGTAGTTGCTCCTCTTC), Synaptophysin (F; CCTGTCCGATGTGAAGATGG, R; AGGTTCAGGAAGCCAAACAC) gene expression. Relative gene expression levels were obtained by normalization to the expression levels of 18S ribosomal RNA (Cdkn1b and Synaptophysin) or β-actin (Mmp9, Mmp2, Nos2, Il-1β).

### Quantitative measurement of islet and pituitary morphology

Islet area, islet nuclei number and pituitary area were measured from hematoxylin and eosin-stained pancreas sections, and Ki67-positive cells were counted from immunohistochemically stained pancreas sections using TissueFAXS (TissueGnostics) and an HS All-in-one Fluorescence microscope BZ-9000 (Keyence).

### Quantification analysis of the HE and IHC images

Image analyses of Insulin, P-S6, P-mTOR, Ki67, and p27 were performed using ImageJ software (imageJ.net). Additionally, color thresholding image adjustment was performed with the Colour Deconvolution plugin (imagej.net/Colour_Deconvolution). We used an optimized immunohistochemistry image processing protocol for DAB staining to semi-quantitatively analyze protein expression, as previously described ^41^. Image analyses of vessels were also done using ImageJ software. Images analyses of Figs. S1 and S4 were analyzed with an HS All-in-one Fluorescence microscope BZ-9000 (KEYENCE).

### Patients analyzed in the study

A retrospective cohort study was conducted consisting of 56 patients who had been pathologically diagnosed with PanNETs and those that had undergone chemotherapy without surgery between July 2014 to August 2021 at National Cancer Center Hospital. Only data from patients who did not receive prior chemotherapy were included, and blood collection was performed before therapy. We did not include patients who had undergone chemotherapy, as these treatments can potentially affect pancreatic function. Some patients had undergone previous surgery and only those who were one year or more post-surgery were included in the analysis. There was no significant difference in the blood glucose levels between the patients with or without surgery (Fig. S6). Chemotherapies included: octreotide, lanreotide, everolimus, sunitinib, streptozotocin, lanreotide+everolimus, lanreotide+sunitinib and others (CDDP+CPT-11, GEM+nabPTX or PRRT). Inclusion criteria were as follows: (i) patients diagnosed with PanNET; (ii) patients with follow-up data; (iii) patients with blood glucose data before the treatments. Cases excluded were (i) patients diagnosed with functional PanNETs or (ii) patients that had developed multiple primary tumors. This study was approved by the Institutional Review Board of the National Cancer Center, Tokyo (approval number; 2013-023). Informed consent was obtained for all cases. Clinical and pathological data were obtained through a detailed retrospective review of the medical records of all patients with PanNET. According to the Standards of Medical Care in Diabetes 2022 issued by the American Diabetes Association ^31^, a fasting blood glucose (FBG) level from 100 to 125 mg/dl is an indication of a prediabetic state. Therefore, high blood glucose levels are defined as blood glucose levels higher than 100 mg/dl. Overall survival (OS) was defined as the length of time (in days) from the beginning of treatment to death from any cause.

### Statistical analysis

Data were calculated and shown as mean ±SEM. Comparisons between the samples were performed by Student *t* test or by one-way ANOVA multiple comparisons (Tukey’s multiple comparisons test) using Prism software (version 6). For the t test, Student’s t test was used when the variances of the groups were not significantly different as evaluated by f test. Survival data were analyzed using Prism software (version 6), and Kaplan–Meyer plots were drawn. Statistical significance was defined as p<0.05. For animal studies, no statistical methods were used for sample size estimate, no samples were excluded, no randomization was used, and no blinding was done.

## ACKNOWLEDGMENTS

We thank Dr. Marc Lamphier for critical reading of the manuscript. Y.C. is a JSPS International Research fellow. This work was carried out by the joint research program of the Institute for Molecular and Cellular Regulation, Gunma University. This study was partly supported by a Grant-in-Aid for Scientific Research (B, #20H03523 to R.O.), Grant-in-Aid for Scientific Research (C, #22K07181 to Yuko T.) and Grant-in-Aid for Young Scientist (B) (#19K16732 to Y.C.) from MEXT of Japan; P-Direct and P-Create from AMED of Japan (R.O.); Grant-in-Aid for JSPS fellow (# 20F20414 to R.O. and Y.C.); research grants of the Okinaka Memorial Institute for Medical Research (R.O.), the Life Science Foundation of Japan (R.O.), Pancreas Research Foundation of Japan (Y.C.) and Ichiro Kanehara Foundation for the promotion of Medical Sciences and Medical Care (Y.C.).

## AUTHOR CONTRIBUTIONS

Conceptualization: RO; Formal Analysis: YC, TY, YuraT, TM, TT, YS, NS, SH, AS, YukoT, RO; Investigation and Methodology: YC, TY, YuraT, TM, TT, YS, NS, AY, SH, AS, YukoT, RO; Funding acquisition: YC, YukoT, RO; Supervision: RO; Writing – original draft; RO, review & editing: YC, TY, YuraT, TM, TT, YS, NS, AY, SH, AS, YukoT, RO.

## Supplementary Figure legends

**Fig. S1 Ketogenic diet suppresses non-functional PanNET development.**

Islet areas of 45-week-old wild-type mice fed a normal diet (N=3), *Men1*^f/f^-RipCre^+^ mice fed a normal diet (N=3) or *Men1*^f/f^-RipCre^+^ mice fed a ketogenic diet from 7-11 weeks of age (N=3) were analyzed. Three pancreatic sections were analyzed for each mouse. The distance between the first and the third section is more than 20 μm. The percentages of islet areas within the total pancreas areas are shown. Data are represented as the mean fold expression ± SD.

**Fig. S2 Ketogenic diet results in lower blood glucose levels.**

Blood glucose levels of 20-week-old male *Men1*^f/f^-RipCre^+^ mice fed a normal diet (N=6) or 40-week-old female *Men1*^f/f^-RipCre^+^ mice fed a ketogenic diet from 10 weeks of age (N=5). Blood glucose levels at basal, after fasting (overnight) and after re-feeding the usual meal for 1.5 hrs were analyzed. Data are represented as the mean fold expression ± SEM.

**Fig. S3 Ketogenic diet does not affect *Cdkn1b* (p27) mRNA expression.**

Islets were isolated from 45-week-old *Men1*^f/f^-RipCre^+^ mice fed a normal diet (N=6) or *Men1*^f/f^-RipCre^+^ mice fed a ketogenic diet from 10 weeks of age (N=5). mRNAs were purified and expression of *Cdkn1b* (p27) was analyzed. Data are represented as the mean fold expression ± SD.

**Fig. S4 Ketogenic diet suppresses non-functional PanNET progression.**

Islet areas of 45-week-old *Men1*^f/f^-RipCre^+^ mice fed a normal diet (N=4) or *Men1*^f/f^-RipCre^+^ mice fed a ketogenic diet from 30 weeks of age (N=4) were analyzed as in Fig. S1.

Three pancreatic sections were analyzed for each mouse. Data are represented as the mean fold expression ± SD.

**Fig. S5 Kaplan–Meier plots of overall survival of the *Men1*^f/f^-RipCre^+^ mice.**

In total, 28 male and 42 female *Men1*^f/f^-RipCre^+^ mice fed a normal diet were analyzed.

**Fig. S6 Blood glucose levels of the patient with or without prior surgery.**

In total, 54 patients with (n=21) or without (n=33) prior surgery were analyzed. Data are represented as the mean fold expression ± SEM.

