## Supplementary material for "Metabolic intervention by low carbohydrate diet suppresses the onset and progression of neuroendocrine tumors": Suppl Figs

### Fig. S1

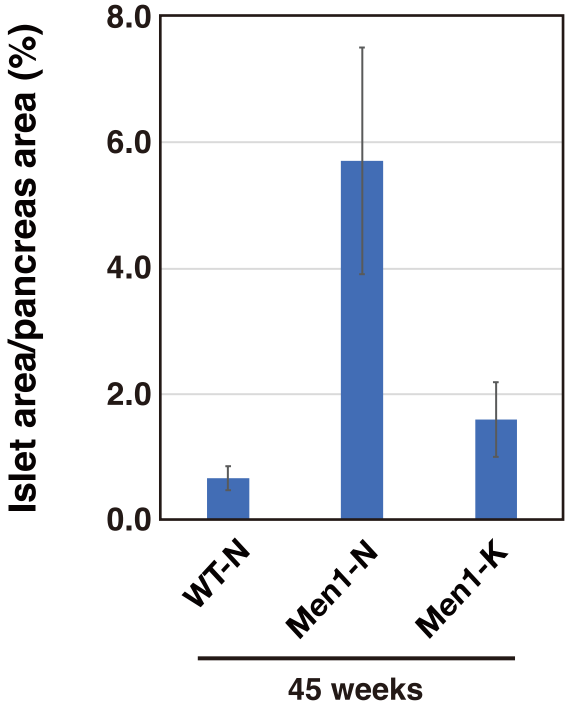


| Tukey's multiple comparisons test | Summary | P Value |
|---|---|---|
| WT-N(45) vs. Men1-N(45) | **** | < 0.0001 |
| WT-N(45) vs. Men1-K-30w(45) | ns | 0.195 |
| Men1-N(45) vs. Men1-K-30w(45) | **** | < 0.0001 |

**Fig. S1 Ketogenic diet suppresses non-functional PanNET development.**

Islet areas of 45-week-old wild-type mice fed a normal diet (N=3), *Men1*$^{f/f}$-RipCre$^{+}$ mice fed a normal diet (N=3) or *Men1*$^{f/f}$-RipCre$^{+}$ mice fed a ketogenic diet from 7-11 weeks of age (N=3) were analyzed. Three pancreatic sections were analyzed for each mouse.

The distance between the first and the third section is more than 20 μm.

The percentages of islet areas within the total pancreas areas are shown.

Data are represented as the mean fold expression ± SD.

**Fig. S2**

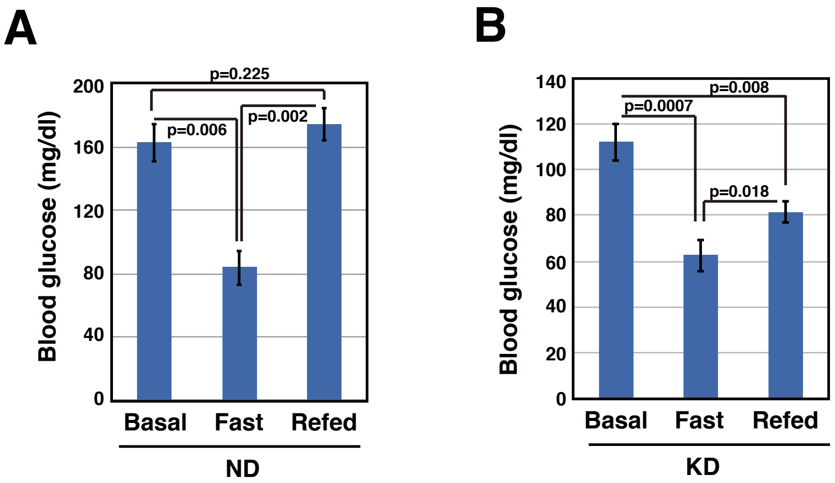


**Fig. S2 Ketogenic diet results in lower blood glucose levels.**

Blood glucose levels of 20-week-old male *Men1*$^{f/f}$-RipCre$^{+}$ mice fed a normal diet (N=6) or 40-week-old female *Men1*$^{f/f}$-RipCre$^{+}$ mice fed a ketogenic diet from 10 weeks of age (N=5). Blood glucose levels at basal, after fasting (overnight) and after re-feeding the usual meal for 1.5 hrs were analyzed. Data are represented as the mean fold expression ± SEM.

### Fig. S3

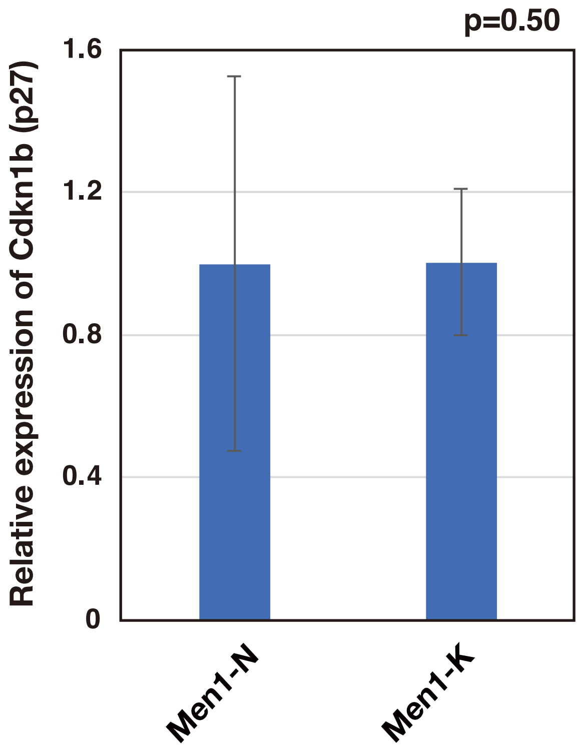


**Fig. S3 Ketogenic diet does not affect *Cdkn1b* (p27) mRNA expression.**

Islets were isolated from 45-week-old *Men1*$^{f/f}$-RipCre$^{+}$ mice fed a normal diet (N=6) or *Men1*$^{f/f}$-RipCre$^{+}$ mice fed a ketogenic diet from 10 weeks of age (N=5).

mRNAs were purified and expression of *Cdkn1b* (p27) was analyzed.

Data are represented as the mean fold expression ± SD.

#### Fig. S4

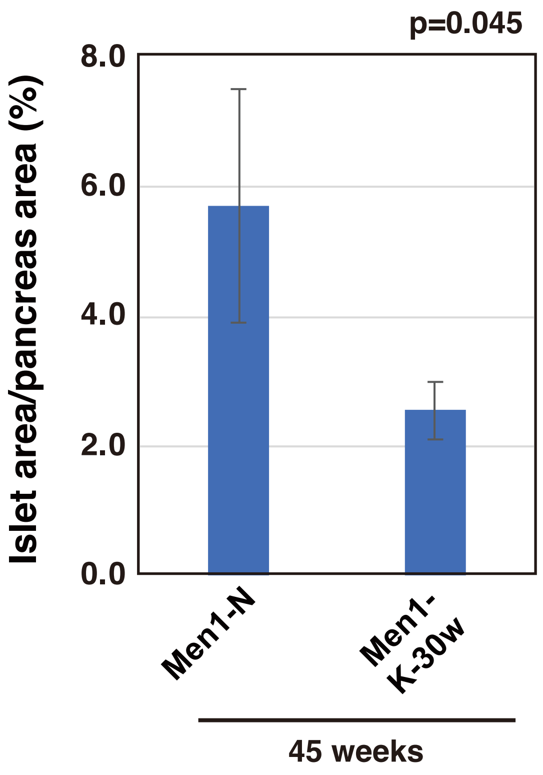


**Fig. S4 Ketogenic diet suppresses non-functional PanNET progression.**

Islet areas of 45-week-old *Men1*$^{f/f}$-RipCre$^{+}$ mice fed a normal diet (N=4) or *Men1*$^{f/f}$-RipCre$^{+}$ mice fed a ketogenic diet from 30 weeks of age (N=4) were analyzed as in Fig. S1.

Three pancreatic sections were analyzed for each mouse.

Data are represented as the mean fold expression ± SD.

### Fig. S5

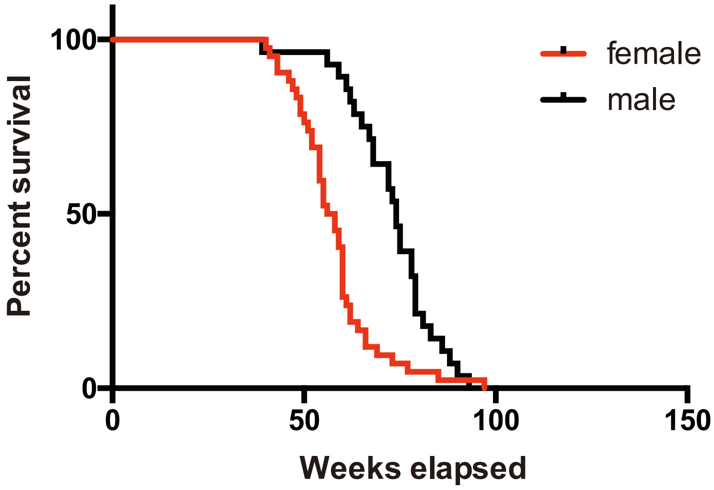


**Fig. S5 Kaplan–Meier plots of overall survival of the *Men1*$^{f/f}$-RipCre$^{+}$ mice.**

In total, 28 male and 42 female *Men1*$^{f/f}$-RipCre$^{+}$ mice fed a normal diet were analyzed.

### Fig. S6

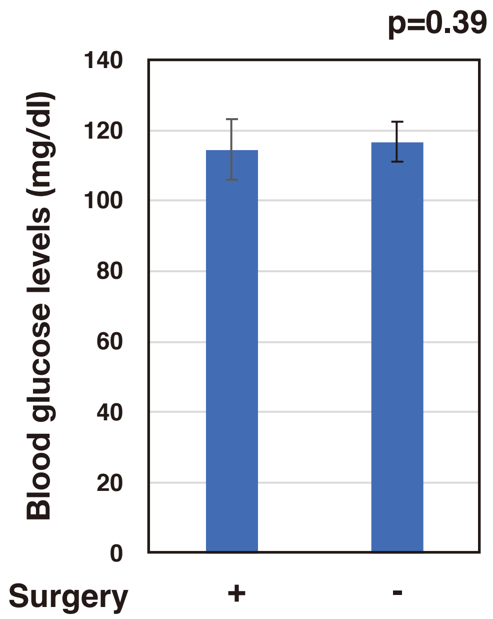


**Fig. S6 Blood glucose levels of the patient with or without prior surgery.**

In total, 54 patients with (n=21) or without (n=33) prior surgery were analyzed.

Data are represented as the mean fold expression ± SEM.
